# Predicted the impacts of climate change and extreme-weather events on the future distribution of fruit bats in Australia

**DOI:** 10.1101/2021.05.13.443960

**Authors:** Vishesh L. Diengdoh, Stefania Ondei, Mark Hunt, Barry W. Brook

**Author notes:** **Author Contributions** VLD proposed and designed the study, analysed the data, and wrote the manuscript. SO contributed significantly to the study design and writing of the manuscript. MH contributed to the writing of the manuscript. BWB contributed significantly to the study design and writing of the manuscript. **Biosketch** VLD is a PhD student at the Dynamics of Eco-Evolutionary Patterns (DEEP) research group at the University of Tasmania, Australia. He is interested in understanding how land-use and land-cover change and climate change impact pollinators. Corresponding Author: Vishesh L. Diengdoh, e. e.

## Abstract

**Aim:** Fruit bats (Megachiroptera) are important pollinators and seed dispersers whose distribution might be affected by climate change and extreme-weather events. We assessed the potential impacts of those changes, particularly more frequent and intense heatwaves, and drought, on the future distribution of fruit bats in Australia. We also focus a case study on Tasmania, the southernmost island state of Australia, which is currently devoid of fruit bats but might serve as a future climate refugium.

**Location:** Australia (continental-scale study) and Tasmania.

**Methods:** Species distribution modelling was used to predict the occurrence of seven species of fruit bats, using an ensemble of machine-learning algorithms. Predictors included extreme-weather events (heatwave and drought), vegetation (as a proxy for habitat) and bioclimatic variables. Predictions were made for the current-day distribution and future (2050 and 2070) scenarios using multiple emission scenarios and global circulation models.

**Results:** Changes in climate and extreme-weather events are forecasted to impact all fruit-bat species, with the loss and gain of suitable areas being predominantly along the periphery of a species’ current distribution. A higher emission scenario resulted in a higher loss of areas for Grey-headed flying fox (*Pteropus poliocephalus*) and Spectacled flying fox (*P. conspicillatus*) but a higher gain of areas for the Northern blossom bat (*Macroglossus minimus*). The Grey-headed flying fox (*Pteropus poliocephalus*) is the only study species predicted to potentially occur in Tasmania under future scenarios.

**Main conclusions:** Fruit bats are likely to respond to climate change and extreme weather by migrating to more suitable areas, including regions not historically inhabited by those species such as Tasmania—possibly leading to human-wildlife conflicts. Conservation strategies (e.g., habitat protection) should focus on areas we found to remain suitable under future scenarios, and not be limited by state-political boundaries.

## Introduction

The projected increase in global temperature and the frequency and intensity of extreme-weather events (IPCC 2014) is a cause of concern for biodiversity protection worldwide, as these changes are observed and predicted to impact the diversity, distribution and mortality of various taxa (McKechnie and Wolf 2010; Newbold 2018; Parmesan and Yohe 2003; Titley et al. 2021; Welbergen et al. 2007). Species are expected to respond to increasing temperatures and extreme weather by shifting their distribution towards more suitable areas (Maxwell et al. 2019; Stillman 2019), likely including new habitats and geopolitical areas (Titley et al. 2021). Range shifts can result in changes in community structure and ecosystem processes which can have positive and negative implications; assessing these changes and implications is essential for developing conservation policies (Wallingford et al. 2020). Fruit bats (Megachiroptera) are an important climate-impact case study. They are a visually striking flying mammal (easily noticed and forming large colonies) that play a critical ecological role in pollination and seed dispersal and can carry large pollen loads and seeds over long distances (Fleming et al. 2009). Their regional diversity and distributional range are expected to be impacted by climate change across several areas within their global extent, which includes Africa, Nepal, Southeast Asia, and Australia (Arumoogum et al. 2019; Hughes et al. 2012; Thapa et al. 2021; Welbergen et al. 2007). Extreme weather events have a particularly strong impact on flying foxes (*Pteropus* spp.), with heatwaves causing many recorded mass-mortality events (Kim and Stephen 2018; Welbergen et al. 2014; Welbergen et al. 2007).

In Australia, correlative species distribution models (SDM) have been used, in a few cases, to assess the impacts of climate change on fruit bats, but either failed to account for changes in extreme weather (see Graham et al. 2019) or did not use species-specific temperature thresholds (see Morán□Ordóñez et al. 2018). The latter is a potentially problematic limitation because temperatures above 42 °C are considered as heatwaves for flying foxes (Ratnayake et al. 2019; Welbergen et al. 2007) while for other species such as bettongs (genus *Bettongia)* temperatures above 28 °C are considered as heatwaves (Bateman et al. 2012). To date, no study has combined projections of future climate with the estimated future frequency and intensity of extreme-weather events (heatwaves and drought) to predict the potential changes in the distribution of fruit bats in Australia—yet this arguably constitutes among the greatest threats to Megachiroptera and bats in general; changes in temperature and precipitation, lack of availability of water, and natural abiotic factors such as weather and fire can impact the survivability and distribution of bats (O’shea et al. 2016; Sherwin et al. 2013).

Fruit bats could respond to climate change and extreme-weather events by shifting their distribution to more suitable areas, because they are highly agile and often travel large distances in search of resources (Roberts et al. 2012; Tidemann and Nelson 2004), with a track record of being able to colonise previously uninhabited areas (Boardman et al. 2020; Parris and Hazell 2005; Westcott and McKeown 2014). In this regard, it is also worth considering whether Tasmania, the southernmost island state of Australia, which historically lacks fruit bats, might be suitable under future climate change. Colonisation by fruit bats to previously uninhabited areas can have significant consequences, as they are a frequent subject of human-wildlife conflicts (Roberts et al. 2012; Tait et al. 2014).

This study aims to assess the potential impacts of climate change and extreme-weather events (heatwaves and drought) on fruit bats in Australia, with particular emphasis on the suitability of Tasmania as a future climate refugium. We used correlative SDMs and an ensemble of machine-learning algorithms (hereafter: algorithms) to estimate the occurrence of fruit bats under current and future climate scenarios (multiple years, emission scenarios and global circulation models). We highlighted which existing geographical areas will become climatically unsuitable and/or new areas suitable for fruit bats in the future and recommend potential strategies to help conserve fruit bats in Australia.

## Methods

### Study species

Australia has 13 species of fruit bats (Hall and Richards 2000). We downloaded records of species occurrence (presence) from Atlas of Living Australia (Atlas of Living Australia 2020). Records were limited to those from 1960 and onwards and classified as ‘human observation’. We removed duplicate records based on latitude and longitude and dubious records (e.g., outliers well outside the known distribution ranges).

We selected those species reported as present in at least 20 different cells of 0.05° spatial resolution (∼ 5 km). Consequently, the Grey-headed flying-fox (*Pteropus poliocephalus*), Little red flying-fox (*P. scapulatus*), Black flying-fox (*P. alecto*), Spectacled flying-fox (*P. conspicillatus*), Common blossom bat (*Syconycteris australis*), Northern blossom bat/Dagger-toothed long-nosed fruit bat (*Macroglossus minimus*), and the Eastern tube-nosed bat (*Nyctimene robinsoni*) were modelled in this study; the other species did not have sufficient data. The IUCN Red List (IUCN 2020) lists only the Grey-headed flying fox as vulnerable (Lunney et al. 2008) and the Spectacled flying fox as endangered (Roberts et al. 2020), while the other species are classified as ‘least concern’.

Presence data was reduced to a single observation per grid cell. Pseudo-absences were generated using the target□group method (Phillips et al. 2009) as it has been shown to robustly handle sampling bias, especially concerning visitation (Fithian et al. 2015). The method consists of generating pseudo-absences for each species by using the presence points of other fruit bats.

### Study area

Fruit bats occur along the coastal margin of the Australian continent, and in some inland regions (Hall and Richards 2000). They are found across a range of habitats, including rainforests, savannas, and urban areas (Hall and Richards 2000; Tait et al. 2014; Westcott 2010). Restricting the geographic extent using a geographical criterion can improve model performance (Acevedo et al. 2012). We examined habitat suitability across the Australian continent, with the exclusion of desert regions, as identified using a Köppen-Geiger climatic layer (Beck et al. 2018), because we assumed it is unlikely that these regions will be suitable for fruit bats due to their high temperatures and extreme range. We also used land-cover data (version 2.1; Lymburner et al. 2015) to mask out land-cover categories we do not expect fruit bats to occupy: extraction sites, salt lakes and alpine grassland.

### Study predictors

To model species occurrence, we selected heatwave, drought, vegetation, and bioclimatic variables as predictors. These predictors all have a spatial resolution of 0.05° (∼ 5 km). Bioclimatic variables and vegetation were resampled using the bilinear interpolation method and nearest neighbour method, respectively, to match the spatial resolution of the heatwave and drought predictors. Resampling was carried out using the *raster* R package (Hijmans 2020).

Heatwave refers to the number of days when maximum or minimum temperature is equal to or greater than 42 °C are tallied in both the historic and projected future 30-year daily time series and averaged annually over the 30-years. Drought refers to the number of months falling below the historic 10^th^ percentile rainfall total were counted in both the historic and projected future 30-year monthly time series and averaged annually over the 30-years.

Heatwaves have caused multiple mortality events in flying foxes (*Pteropus* spp.; Kim and Stephen 2018; Welbergen et al. 2014; Welbergen et al. 2007). We are unaware of similar events being recorded in other, rarer (less conspicuous) fruit-bat species– *Nyctimene* spp., *Macroglossus* spp., and *Syconycteris* spp., however, we assumed these fruit bat genera to have the same temperature threshold as flying foxes (*Pteropus* spp.). Drought can impact the availability of foraging resources (Lučan et al. 2016) which in turn could influence the occurrence of fruit bats. Heatwave and drought variables were downloaded from Climate Change in Australia (https://www.climatechangeinaustralia.gov.au; Clarke et al. 2011; Commonwealth Scientific and Industrial Research Organisation (CSIRO) and Bureau of Meteorology 2020; Whetton et al. 2012). Given the influence of heatwaves and drought on fruit bats, we selected bioclimatic variables that were a direct measure of temperature and precipitation: minimum, maximum, and mean temperature and precipitation of different months and quarters (see Appendix S1 Table S1.1 in Supporting Information). Bioclimatic variables were downloaded from WorldClim (version 2.1; https://www.worldclim.org; Fick and Hijmans 2017).

Vegetation is important in the selection of foraging and roosting sites for fruit bats (Tidemann et al. 1999). We assumed vegetation to be a constant predictor under current and future scenarios due to lack of data on its future distribution, although it is likely to eventually track climate shifts too. Vegetation data (major groups) were downloaded from the National Vegetation Information System (version 5.1; Australian Government Department of Agriculture Water and the Environment 2018). These data consisted of thirty-three vegetation classes, which were reduced to fourteen macro classes (see Appendix S1 Table S1.2).

To account for the influence of climate models and emission scenarios on SDMs (Brun et al. 2020; Goberville et al. 2015; Thuiller et al. 2019), we selected multiple global circulation models (GCMs) and representative concentration pathways (RCP) to model species occurrence for 2050 and 2070. The bioclimatic and extreme-weather-events data sources had different GCMs available, limiting our choice to GCMs common across the two data sources.

This left three GCMs: Australian Community Climate and Earth System Simulator 1.0 (ACCESS 1.0; Bi et al. 2013), Hadley Centre Global Environment Model Carbon Cycle (HadGEM2-CC; Collins et al. 2011), and the Model for Interdisciplinary Research on Climate (MIROC5; Watanabe et al. 2010). We selected RCP 4.5 (Thomson et al. 2011) and 8.5 (Riahi et al. 2011) because they represented medium- and high-emission scenarios respectively, and best match the current emissions trajectory. The bioclimatic variables from the multiple GCMs and RCPs were downloaded from WorldClim (http://worldclim.com/; version 1.4; Hijmans et al. 2005).

### Training data

The training data were biased to pseudo-absences with presence to pseudo-absence ratio set to a maximum of 1:10. Pseudo-absences points were selected using a stratified random method to ensure an adequate number of vegetation class (the only factorial predictor) were represented, and thereby result in better training data. The training data for each species will be made public via figshare with the following DOI https://doi.org/10.6084/m9.figshare.14583282. However, currently, the data can be shared privately on request.

### Variable selection

We reduced the number of predictors by removing correlated variables using a threshold value of 0.7 (Dormann et al. 2013). We used the *findCorrelation* function in the *caret* R package (Kuhn 2020) to remove, for each pair of highly correlated variables, the one with the largest mean absolute correlation. Instead of using all the non-correlated predictors to fit the models, we created a list of all possible combinations of three non-correlated bioclimatic predictors and included heatwave, drought, and vegetation predictors since they were not strongly correlated to each other or any of the climate variables. We constrained any given approach to a maximum of six predictors, to keep the models relatively simple. This resulted in a set of four different groups of predictors/candidate models for each species (see Appendix S1 Table S1.3).

### Cross-validation

We used repeated 70/30% training/test cross-validation splits with 50 repeats for algorithm tuning/optimisation, selecting predictor/candidate models for each species, and calculating the variable importance. Accuracy was evaluated using Area Under the Receiver Operating Characteristic Curve (AUC) and True Skill Statistic (TSS) (Allouche et al. 2006) of the test (hold-out) data. The algorithms used in the study include random forest (RF), classification and regression trees (CART), neural network (NN), stochastic gradient boosting (GB), penalised generalized linear model (GLM), penalised multinomial regression (MR) and flexible discriminant analysis (FDA). These algorithms were selected as they are commonly available in species distribution modelling R packages (see Naimi and Araújo 2016; Schmitt et al. 2017). Cross-validation was implemented using the unified *caret* and *caretEnsemble* R packages (Deane-Mayer and Knowles 2019; Kuhn 2020). The accuracy metrics achieved by the predictor/candidate models are provided in the supplementary material (see Appendix S1 Table S1.3).

### Species occurrence models

Occurrence models were then re-fitted to the full dataset using the best predictor/candidate models derived from the cross-validation step. We also assessed the partial dependence plots of each algorithm and then averaged them (unweighted) using the *pdp* r package (Greenwell 2017). Given that SDMs are influenced by algorithms and climate models (Brun et al. 2020; Thuiller et al. 2019) we selected an ensemble modelling method. An ensemble algorithm averages the prediction of structurally different algorithms and could potentially overcome uncertainty in model selection and improve prediction accuracy by reducing variance and bias (Dormann et al. 2018). As such, we averaged the algorithms using an unweighted method and assessed the AUC and TSS (Allouche et al. 2006) of the final ensemble model. Future-occurrence predictions were made for each species per year per emission scenario per climate model and then ensembled (unweighted) per year per emission scenario for each species.

### Species presence-absence models

We used the sensitivity□specificity sum maximisation approach (Liu et al. 2005) to select the optimal suitability threshold for transforming the probabilistic ensemble occurrence models to binary (presence and absence representations). Using the prediction under current climatic conditions as a reference, we determined which areas (i.e., number of occupied grid cells) will be stable, lost, or gained under future climatic scenarios. Stable areas are those that remain climatically suitable or unsuitable under current and future scenarios. Lost areas are those currently suitable but predicted to become unsuitable under future scenarios. Gained areas are those that are currently unsuitable but predicted to become suitable under future scenarios. We additionally assessed the presence of a species under different climate types. We assess the presence of a species under current conditions and future scenarios using the present and future climate type by Beck et al. (2018), respectively.

The study method is additionally described using the Overview, Data, Model, Assessment and Prediction (ODMAP) protocol (Zurell et al. 2020; see Appendix S2). All the analyses were done using R software (R Core Team 2020).

## Results

The ensemble models for the Grey-headed flying fox, Spectacled flying fox, and Northern blossom bat under current climatic conditions achieved high descriptive accuracy (Table 1; Fig.1). The models for the Little red flying fox, Black flying fox, Eastern tube-nosed, and Common blossom bat achieved lower accuracy metrics (AUC 0.7-0.8 and TSS 0.2-0.4) during cross-validation, and as a result, we did not predict the occurrence for these species (see Appendix S1 Table S1.3). Predictions for climate-change scenarios were thus made for only three species: Grey-headed flying fox, Spectacled flying fox, and Northern blossom bat (Fig. 2). The predicted ensemble occurrence models for the Grey-headed flying fox, Spectacled flying fox, and Northern blossom bat will be made publicly available as GeoTiff images via https://doi.org/10.6084/m9.figshare.14583282.

**Table 1.** Average cross-validated AUC and TSS values attained by the best candidate model for each species along with their ensemble goodness of fit AUC and TSS values. N.B. the average CV and ensemble GOF metrics are not comparable to each other. The CV metrics were derived from the test data during cross-validation while the GOF metrics were derived using the entire training data.

| Species | Predictors | Average<br>CV<br>AUC | Average<br>CV<br>TSS | Ensemble<br>GOF<br>AUC | Ensemble<br>GOF<br>TSS |
| --- | --- | --- | --- | --- | --- |
| Grey-headed flying fox | drt + hwave + veg + bio19 + bio8 + bio9 | 0.896 | 0.640 | 0.833 | 0.666 |
| Spectacled flying fox | drt + hwave + veg + bio18 + bio17 + bio8 | 0.996 | 0.965 | 0.997 | 0.995 |
| Northern blossom bat | drt + hwave + veg + bio18 + bio5 + bio9 | 0.941 | 0.660 | 0.964 | 0.929 |

\* *drt* = drought, *hwave* = heatwave, *veg* = vegetation, *bio5* = max temperature of warmest month, *bio8* = mean temperature of wettest quarter, *bio9* = mean temperature of driest quarter, *bio18* = precipitation of warmest quarter, *bio17* = precipitation of driest quarter, *bio19* = precipitation of coldest quarter.

**Figure 1.**
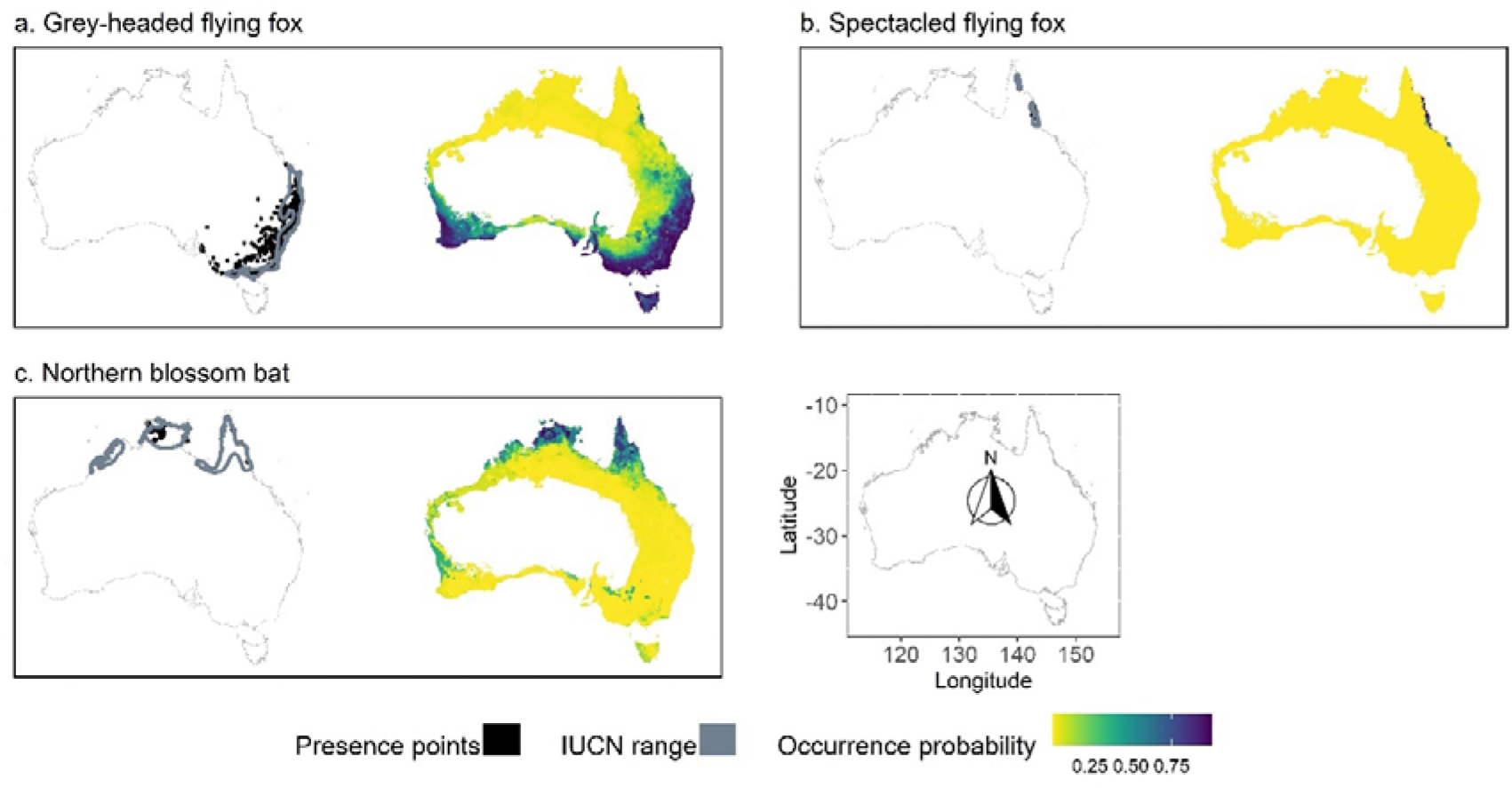
Predicted probability of occurrence of the (a) Grey-headed flying-fox, (b) Spectacled flying-fox, and (c) Northern blossom bat under current climatic conditions along with their presence points and IUCN range boundaries.

**Figure 2.**
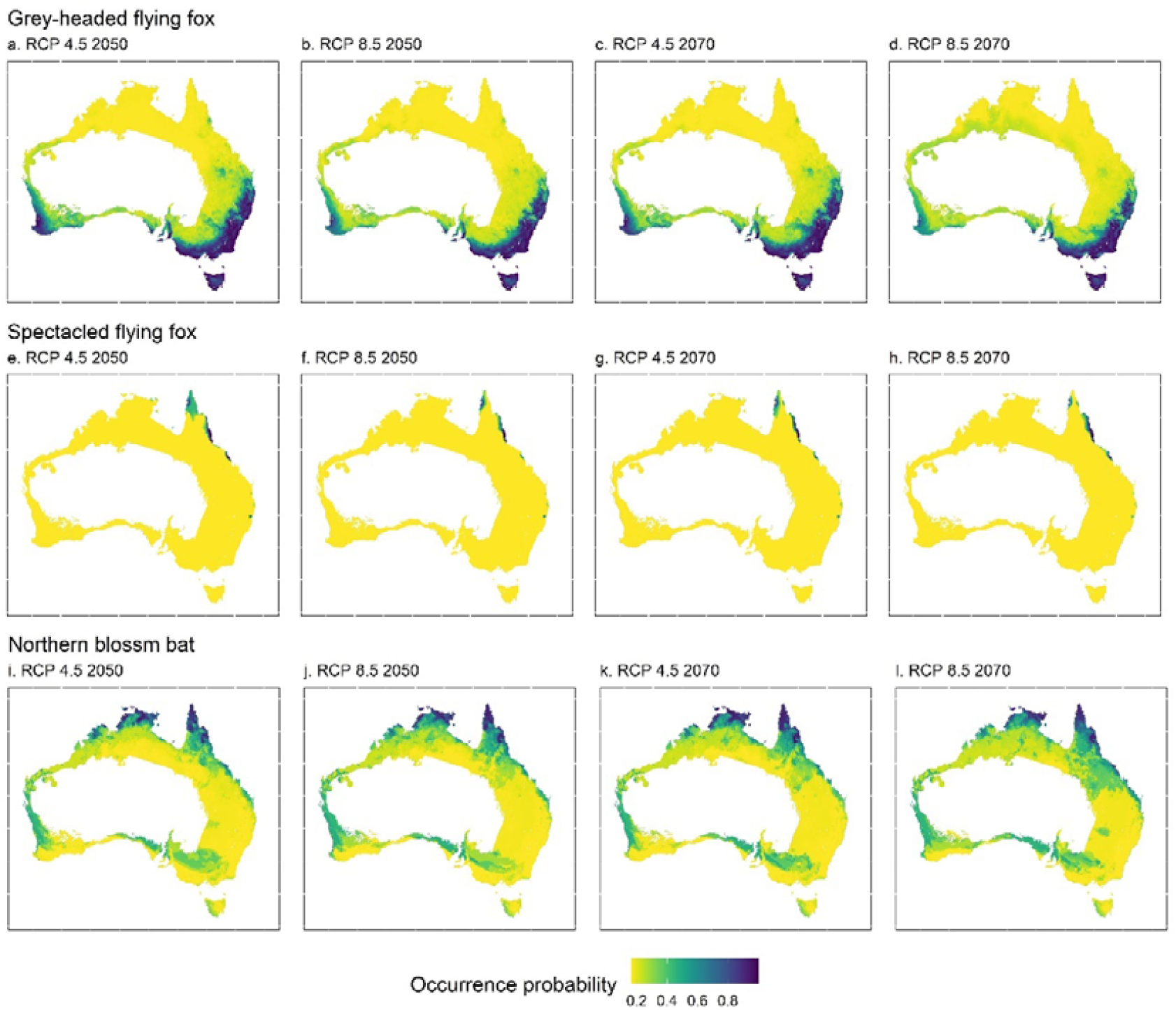
Predicted probability of occurrence for the Grey-headed flying-fox (a-d), Spectacled flying-fox (e-h), and Northern blossom bat (i-l) under different future climatic scenarios.

Overall, mean temperature of wettest quarter, precipitation of warmest quarter, and mean temperature of driest quarter, were the most important predictors for the Grey-headed flying fox, Spectacled flying fox, and Northern blossom bat models, respectively (see Appendix S1 Table S1.4). The extreme weather-event predictors (heatwave and drought) were among the top five important variables for all the models. The vegetation class of non-native vegetation was the most important for the Grey-headed flying fox and Spectacled flying fox, while the vegetation class Eucalypt was the most important for the Northern blossom bat (see Appendix S1 Table S1.4).

Based on the partial-dependence plots we found, the probability of occurrence of the Grey-headed flying fox decreases with the increasing number of days of drought (range from 0.5 to 1.4 days). However, its occurrence increases with the number of days of heatwaves (0.0 to 0.4 days) up to 0.3 days after which it decreases (see Appendix S1 Fig. S1.1). For the Spectacled flying fox, the occurrence decreases with the increasing number of days of drought (0.5 to 1.4 days) and heatwave (0.0 to 0.5 days; see Appendix S1 Fig. S1.2) while for the Northern blossom bat, the occurrence increases and decreases with the increasing number of days of drought (0.5 to 1.3 days) and heatwave (0.0 to 0.5 days), respectively (see Appendix S1 Fig. S1.3). The vegetation types of Acacia, Mallee, and Rainforest were associated with a lower probability of occurrence of the Grey-headed flying fox, while no major differences were detected among the other vegetation types (Appendix S1 Fig. S1.1). Eucalypt and non-native vegetation types have the highest positive influence on the Spectacled flying fox, which has a low probability of occurrence in all the other vegetation types (Appendix S1 Fig. S1.2). Eucalypt, rainforest and other shrublands have the highest positive influence on the Northern blossom bat, while Acacia and Grassland vegetation are predicted to have the least influence (Appendix S1 Fig. S1.3).

The severity of impacts of climate change and extreme weather events varied between species, and so did the extent of areas lost and gained, which depended on the year and emission scenario considered (Fig. 3). As expected, areas lost and gained were, overall, predicted to occur along the edges of a species’ current distribution range (Fig. 3). For the Grey-headed flying fox and Spectacled flying fox, the percentage of areas lost was predicted to be higher in the year 2070 than 2050 and higher under 8.5 than 4.5 emission scenario. Conversely, for the Northern blossom bat, the percentage of areas lost was predicted to be higher in the year 2050 than 2070 and higher under 4.5 than 8.5 emission scenario (see Appendix S1 Table S1.5). For the Grey-headed flying fox and Northern blossom bat, the percentage of areas gained was forecasted to be higher in the year 2070 than 2050 and under 8.5 than 4.5 emission scenario. However, for the Spectacled flying fox, the percentage of areas gained was predicted to be higher in the year 2050 than 2070 and higher under 4.5 than 8.5 emission scenario (Appendix S1 Table S1.5). The binary (presence and absence) models for the Grey-headed flying fox, Spectacled flying fox, and Northern blossom bat will be made publicly available as GeoTiff images via https://doi.org/10.6084/m9.figshare.14583282.

**Figure 3.**
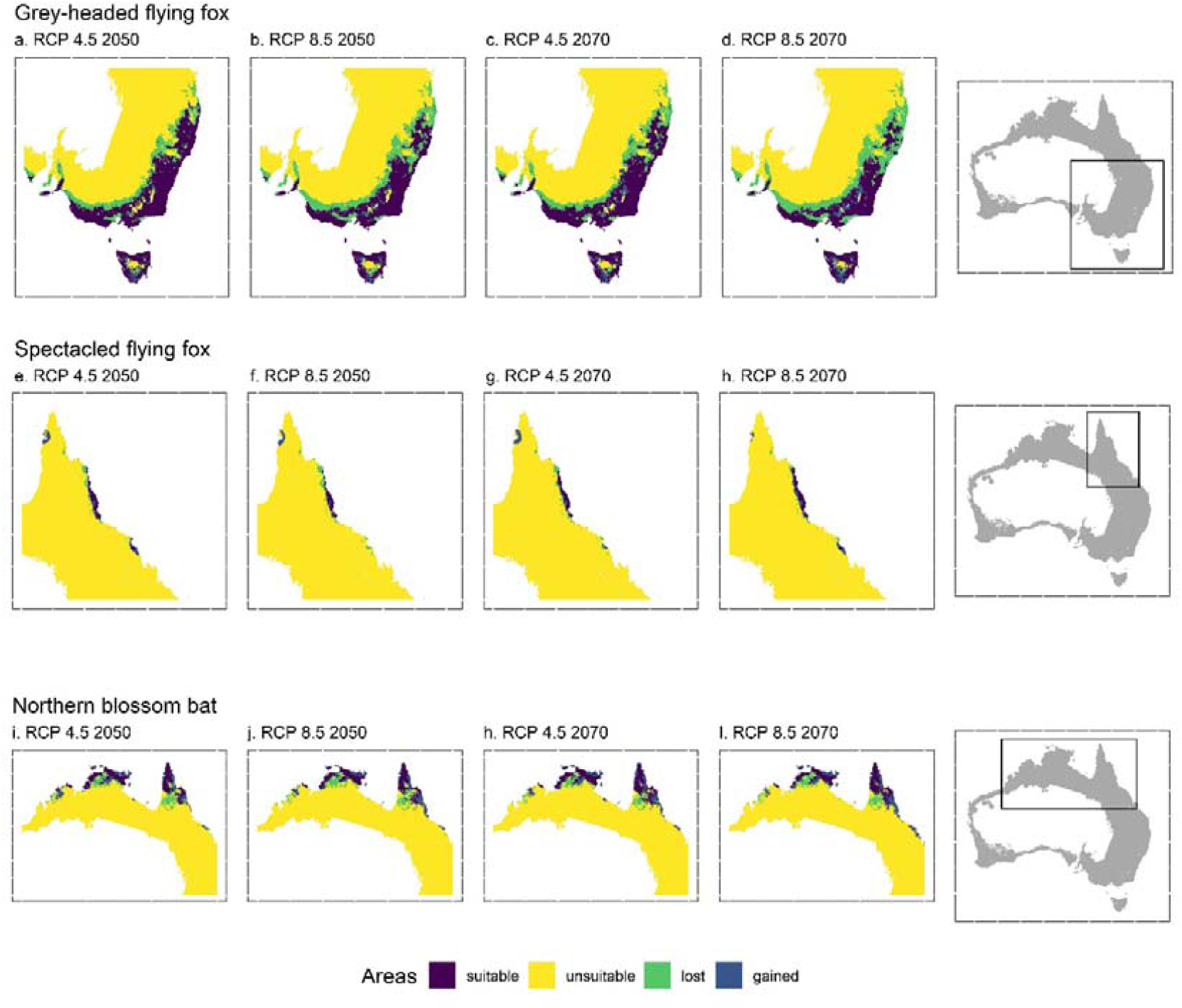
Predicted area suitability for the Grey-headed flying-fox (a-d), Spectacled flying-fox (e-h), and Northern blossom bat (i-l) under different future climatic scenarios.

Under current conditions, the occurrence of the Grey-headed flying fox was found to be highest in temperate no dry season warm-summer climate and temperate no dry season hot summer climate which are found along the southeast of Australia including Tasmania (see Appendix S1 Table S1.6; climate descriptions are based on Köppen-Geiger climate classification, see Beck et al. 2018). Under future scenarios, the occurrence of the Grey-headed flying fox increases in temperate no dry season hot summer climate but decreases in temperate dry summer warm summer and arid steppe cold climate (see Appendix S1 Table S1.6). The occurrence of the Spectacled flying fox is highest in tropical and temperate dry winter hot summer climate under current conditions. These climates are found in the north and northeast of Australia. However, this species is not predicted to occur in temperate dry winter hot summer climate under future scenarios (see Appendix S1 Table S1.6). The occurrence of the Northern blossom bat is highest in tropical savannah climate under current conditions and future scenarios (see Appendix S1 Table S1.6). Tropical savanna is found in northern Australia.

Of the species studied, only the Grey-headed flying fox is predicted to occur in Tasmania and under current conditions and future scenario (Fig. 2a-d). The Grey-headed flying fox is also predicted to occur across the southwest of Australia, which is also devoid of fruit bats, under current conditions. However, unlike Tasmania, the occurrence in southwest Australia is predicted to decrease with increasing emission scenario and year (Fig. 2a-d).

## Discussion

Fruit bats are important vertebrate pollinators (Fleming et al. 2009) but are threatened by climate change and extreme weather events (O’shea et al. 2016; Sherwin et al. 2013). We assessed the impacts of those factors on the occurrence and abundance of fruit bats in Australia using correlative SDMs with an emphasis on Tasmania as a potential future refugium under climate change. Of the seven species studied, we obtained reliable models (under current conditions) for only the Grey-headed flying fox, Spectacled flying fox, and Northern blossom bat, and consequently, future distribution was modelled for these three species. We found that the predicted impacts varied between species, year, and emission scenarios in complex ways. Of the three species with forecasts, the Grey-headed flying fox, the only one currently found in temperate areas, was predicted to have suitable conditions in Tasmania under both current and future scenarios.

The overall loss and gain of areas along the edges of species distributional ranges are, in large part, driven by the predicted expansion of hotter climate types into cooler climates in Australia (see Beck et al. 2018). In general, populations along range boundaries are likely to be more sensitive to climate change and extreme weather than those within the core (Parmesan et al. 2000). Tidemann and Nelson (2004) have suggested that the northern edge of the Grey-headed flying fox has already contracted south due to climate change and our results indicate that climate change and extreme weather events will further exacerbate this process. We were unable to find comparative studies for our findings of the Spectacled flying fox and Northern blossom bat.

The Grey-headed flying fox is found in warm temperate to tropical climates (Parris and Hazell 2005), while the Spectacled flying fox is found in the wet tropics (Tait et al. 2014). This could explain why temperate dry summer climate types and temperate hot summer climate are found to be associated with a decrease in suitability for the Grey-headed flying fox and Spectacled flying fox, respectively. The biogeographic region that a bat species occupies can influence its response to climate change (Rebelo et al. 2010).

Our predictions indicate that the Grey-headed flying fox could be supported by the climatic conditions that exist today in Tasmania, even though they are currently not found on the island. Tasmania’s glacial history might have limited the availability of suitable habitat in the past, preventing flying foxes from establishing permanent colonies on the island (Driessen et al. 2011). However, as species can shift their distribution as a response to climate change (Parmesan and Yohe 2003; Titley et al. 2021), particularly endothermic species, such as bats, that track their niche (Araújo et al. 2013), it is plausible that future climate and extreme weather events and the reduction of suitable habitat in their original distributional range could drive the Grey-headed flying fox to Tasmania, despite the 250 km wide oceanic barrier of Bass Strait. Indeed, apparent vagrants have occasionally been recorded on the Bass Strait islands (Driessen 2010).

Shifting the probability density or total range of a species’ distribution as a response to climate change is possible when gradual changes in ambient temperature allow for adjustment of their physiology and behaviour to new conditions. However, this might not be feasible for extreme-weather events such as heatwaves (Bondarenco et al. 2014). This may result in a situation where flying foxes need human-assisted migration to established in places such as Tasmania, to safeguard their population viability. This, in turn, gets to the deeper social question of whether the general public would be willing to tolerate climate-driven ‘refugees’, a term applicable to animals as well (Derham and Mathews 2020), colonising new habitats—such as Tasmania—naturally or through human assistance.

The management of species moving into new geopolitical areas will likely depend on their ecological and socio-economic values (Scheffers and Pecl 2019). Fruit bats can contribute to pollination (a key ecosystem service) in Tasmania and as nocturnal mammal species, they are unlikely to compete with diurnal nectarivorous and frugivorous birds, although they could reduce resource availability (Westcott and McKeown 2014). However, bats are known carriers of diseases, e.g., Hendra virus (Martin et al. 2018), and as such have been persecuted (Hughes et al. 2007; MacFarlane and Rocha 2020) and are a frequent subject of human-wildlife conflicts (Roberts et al. 2012; Tait et al. 2014). This would make their management in new locations a complex and challenging task. In this context, we advocate that, whilst *in*-*situ* protection should be the primary aim of all conservation efforts, there remains a need to speculate on, model, and discuss potential *ex*-*situ* conservation strategies. Our results underscore the importance of implementing conservation strategies focused on areas and climatic zones commonly suitable under all predicted scenarios (Fig. 3), which do not respect geopolitical boundaries.

The unreliable models that resulted from the available occurrences for the Little red flying fox, Black flying fox, Eastern tube-nosed and Common blossom bat were probably due to a combination of sampling inadequacy and/or bias, biotic interactions, and spatial scale. The number and location of presence points can influence model accuracy (van Proosdij et al. 2016; Wisz et al. 2008), limiting the ability to model the distribution of species with few presence points. Occurrence records could also be out of equilibrium with their current environment due to recent past human impacts, such as land-use and land-cover change, which can also degrade model outcomes (Dormann 2007). Finally, the spatial scale of the data used was ∼5 km, which is relatively coarse and might fail to represent important habitat characteristics for some species, as sensitivity to scale can depend on the species attributes (Dormann 2007).

## Conclusion

Our study found climate change and associated changes in extreme-weather events to have generally detrimental, but different impact on fruit-bat species. Although no fruit bats are currently found in the southernmost part of Australia (Tasmania), this large island is found to be climatically suitable for the Grey-headed flying fox, now and in the future. Fruit bats could respond to climate change and extreme-weather events by migrating to more suitable areas or do so via human-assisted migration, to safeguard their population viability. Both instances raise challenging socio-political questions that would benefit from discussion and debate now, rather than at some future crisis point. Although our study assessed the impacts of climate change and extreme weather events of fruit bats in Australia, we are unaware of how these changes would affect the Spectacled flying fox and the Northern blossom bat which are also found in Southeast Asia and on Pacific Islands (Francis et al. 2008; Roberts et al. 2020). Future studies should consider assessing the impacts of climate change and extreme weather events across a species entire distribution range. Future studies should also consider collecting additional field information to obtain more reliable models, mapping movement patterns for fruit bats under climate change and developing mechanistic (physiology-based) SDMs to further improve the focus of conservation efforts.

## Data Accessibility Statement

The training data (csv files) and predicted models (GeoTiff files) will be made publicly available via figshare with the following DOI: https://doi.org/10.6084/m9.figshare.14583282. However, currently, the dataset can be shared privately on request.

## Acknowledgements

We thank John Clarke and Vanessa Round from Climate Change in Australia (https://www.climatechangeinaustralia.gov.au/)/ Commonwealth Scientific and Industrial Research Organisation (CSIRO) for providing the data on extreme weather events. This work was supported by the Australian Research Council [grant number FL160100101].

