## Appendix S1 for "Predicted the impacts of climate change and extreme-weather events on the future distribution of fruit bats in Australia"

**Table S1.1** Bioclimatic variables used in the study.

**Bioclimatic variables**

bio5 = max temperature of warmest month

bio6 = min temperature of coldest month

bio8 = mean temperature of wettest quarter

bio9 = mean temperature of driest quarter

bio10 = mean temperature of warmest quarter

bio11 = mean temperature of coldest quarter

bio13 = precipitation of wettest month

bio14 = precipitation of driest month

bio16 = precipitation of wettest quarter

bio17 = precipitation of driest quarter

bio18 = precipitation of warmest quarter

bio19 = precipitation of coldest quarter

**Table S1.2** Reclassified code of vegetation types.

| **Original code** | **Vegetation Type** | **Reclassified code** |
| --- | --- | --- |
| 6 | Acacia Forests and Woodlands | 6 |
| 16 | Acacia Shrublands | 6 |
| 13 | Acacia Open Woodlands | 6 |
| 25 | Cleared, Non-Native Vegetation, Buildings | 25 |
| 2 | Eucalypt Tall Open Forests | 2 |
| 3 | Eucalypt Open Forests | 2 |
| 4 | Eucalypt Low Open Forests | 2 |
| 12 | Tropical Eucalypt Woodlands/Grasslands | 2 |
| 5 | Eucalypt Woodlands | 2 |
| 11 | Eucalypt Open Woodlands | 2 |
| 19 | Tussock Grasslands | 19 |
| 20 | Hummock Grasslands | 19 |
| 21 | Other Grasslands, Herblands, Sedgelands and Rushlands | 19 |
| 18 | Heathlands | 18 |
| 23 | Mangroves | 24 |
| 24 | Inland Aquatic - freshwater, salt lakes, lagoons | 24 |
| 14 | Mallee Woodlands and Shrublands | 14 |
| 32 | Mallee Open Woodlands and Sparse Mallee Shrublands | 14 |
| 27 | Naturally Bare - sand, rock, claypan, mudflat | 27 |
| 15 | Low Closed Forests and Tall Closed Shrublands | 15 |
| 30 | Unclassified Forest | 7 |
| 7 | Callitris Forests and Woodlands | 7 |
| 8 | Casuarina Forests and Woodlands | 7 |
| 9 | Melaleuca Forests and Woodlands | 7 |
| 10 | Other Forests and Woodlands | 7 |
| 31 | Other Open Woodlands | 7 |
| 26 | Unclassified Native Vegetation | 26 |
| 29 | Regrowth, Modified Native Vegetation | 26 |
| 17 | Other Shrublands | 17 |
| 22 | Chenopod Shrublands, Samphire Shrublands and Forblands | 17 |
| 1 | Rainforests and Vine Thickets | 1 |
| 28 | Sea and Estuaries | 28 |
| 99 | Unknown/No Data | 99 |

**Table S1.3** Cross-validated AUC and TSS scores of the different machine learning algorithms and their average for each predictors/candidate models. Bold predictors/values indicate those selected for model fitting.

| **Species** | **Predictors/candidate models*** | **Accuracy metric**** | **Machine learning algorithm***** | | | | | | | | | | | | | | |  |
| --- | --- | --- | --- | --- | --- | --- | --- | --- | --- | --- | --- | --- | --- | --- | --- | --- | --- | --- |
|  |  |  | **RF** | | **CART** | | **NN** | | **GBM** | | **GLM** | | **MR** | | **FDA** | | **average** | |
| Grey-headed flying fox  (*Pteropus poliocephalus*) | drt + hwave + veg + bio18 + bio19 + bio8 | AUC | 0.921 | | 0.863 | | 0.901 | |  | | 0.886 | | 0.887 | | 0.920 | | 0.897 | |
|  | drt + hwave + veg + bio18 + bio19 + bio9 |  | 0.913 | | 0.854 | | 0.898 | |  | | 0.887 | | 0.887 | | 0.903 | | 0.890 | |
|  | drt + hwave + veg + bio18 + bio8 + bio9 |  | 0.917 | | 0.857 | | 0.891 | |  | | 0.862 | | 0.863 | | 0.911 | | 0.883 | |
|  | **drt + hwave + veg + bio19 + bio8 + bio9** |  | **0.920** | | **0.866** | | **0.907** | |  | | **0.885** | | **0.885** | | **0.917** | | **0.897** | |
|  | drt + hwave + veg + bio18 + bio19 + bio8 | TSS | 0.662 | | 0.643 | | 0.640 | |  | | 0.615 | | 0.616 | | 0.637 | | 0.636 | |
|  | drt + hwave + veg + bio18 + bio19 + bio9 |  | 0.650 | | 0.640 | | 0.636 | |  | | 0.628 | | 0.629 | | 0.636 | | 0.636 | |
|  | drt + hwave + veg + bio18 + bio8 + bio9 |  | 0.666 | | 0.655 | | 0.640 | |  | | 0.622 | | 0.614 | | 0.646 | | 0.640 | |
|  | **drt + hwave + veg + bio19 + bio8 + bio9** |  | **0.661** | | **0.646** | | **0.643** | |  | | **0.627** | | **0.625** | | **0.641** | | **0.641** | |
| Little red flying fox  (*Pteropus scapulatus*) | drt + hwave + veg + bio18 + bio19 + bio8 | AUC | 0.762 | | 0.662 | | 0.713 | |  | | 0.708 | | 0.705 | | 0.728 | | 0.713 | |
|  | drt + hwave + veg + bio18 +bio19 + bio9 |  | 0.756 | | 0.661 | | 0.714 | |  | | 0.702 | | 0.699 | | 0.736 | | 0.711 | |
|  | drt + hwave + veg + bio18 + bio8 + bio9 |  | 0.772 | | 0.671 | | 0.727 | |  | | 0.709 | | 0.702 | | 0.740 | | 0.720 | |
|  | drt + hwave + veg + bio19 + bio8 + bio9 |  | 0.762 | | 0.665 | | 0.724 | |  | | 0.705 | | 0.704 | | 0.730 | | 0.715 | |
|  | drt + hwave + veg + bio18 + bio19 + bio8 | TSS | 0.309 | | 0.259 | | 0.273 | |  | | 0.124 | | 0.231 | | 0.277 | | 0.245 | |
|  | drt + hwave + veg + bio18 +bio19 + bio9 |  | 0.301 | | 0.245 | | 0.266 | |  | | 0.134 | | 0.238 | | 0.250 | | 0.239 | |
|  | drt + hwave + veg + bio18 + bio8 + bio9 |  | 0.326 | | 0.269 | | 0.276 | |  | | 0.114 | | 0.228 | | 0.263 | | 0.246 | |
|  | drt + hwave + veg + bio19 + bio8 + bio9 |  | 0.312 | | 0.269 | | 0.273 | |  | | 0.171 | | 0.225 | | 0.258 | | 0.251 | |
| Black flying fox  (*Pteropus alecto*) | drt + hwave + veg + bio18 + bio19 + bio8 | AUC | 0.865 | | 0.809 | | 0.849 | | 0.863 | | 0.845 | | 0.845 | | 0.858 | | 0.848 | |
|  | drt + hwave + veg + bio18 + bio19 + bio9 |  | 0.868 | | 0.814 | | 0.850 | | 0.873 | | 0.845 | | 0.843 | | 0.867 | | 0.851 | |
|  | drt + hwave + veg + bio18 + bio8 + bio9 |  | 0.867 | | 0.821 | | 0.856 | | 0.870 | | 0.849 | | 0.847 | | 0.868 | | 0.854 | |
|  | drt + hwave + veg + bio19 + bio8 + bio9 |  | 0.861 | | 0.816 | | 0.839 | | 0.865 | | 0.845 | | 0.845 | | 0.861 | | 0.847 | |
|  | drt + hwave + veg + bio18 + bio19 + bio8 | TSS | 0.483 | | 0.436 | | 0.446 | | 0.456 | | 0.388 | | 0.405 | | 0.470 | | 0.441 | |
|  | drt + hwave + veg + bio18 + bio19 + bio9 |  | 0.486 | | 0.473 | | 0.464 | | 0.488 | | 0.367 | | 0.397 | | 0.493 | | 0.453 | |
|  | drt + hwave + veg + bio18 + bio8 + bio9 |  | 0.486 | | 0.454 | | 0.484 | | 0.482 | | 0.238 | | 0.444 | | 0.500 | | 0.441 | |
|  | drt + hwave + veg + bio19 + bio8 + bio9 |  | 0.486 | | 0.450 | | 0.452 | | 0.479 | | 0.199 | | 0.427 | | 0.495 | | 0.427 | |
| Spectacled flying fox  (*Pteropus conspicillatus*) | **drt + hwave + veg + bio18 + bio17 + bio8** | AUC | **0.997** | | **0.998** | | **0.997** | |  | |  | | **0.993** | |  | | **0.996** | |
|  | drt + hwave + veg + bio18 + bio17 + bio9 |  | 0.997 | | 0.998 | | 0.993 | |  | |  | | 0.989 | |  | | 0.994 | |
|  | drt + hwave + veg + bio18 + bio8 + bio9 |  | 0.998 | | 0.998 | | 0.998 | |  | |  | | 0.998 | |  | | 0.998 | |
|  | drt + hwave + veg + bio17 + bio8 + bio9 |  | 0.994 | | 0.902 | | 0.954 | |  | |  | | 0.959 | |  | | 0.952 | |
|  | **drt + hwave + veg + bio18 + bio17 + bio8** | TSS | **0.981** | | **0.996** | | **0.949** | |  | |  | | **0.939** | |  | | **0.966** | |
|  | drt + hwave + veg + bio18 + bio17 + bio9 |  | 0.974 | | 0.995 | | 0.935 | |  | |  | | 0.933 | |  | | 0.959 | |
|  | drt + hwave + veg + bio18 + bio8 + bio9 |  | 0.968 | | 0.996 | | 0.918 | |  | |  | | 0.904 | |  | | 0.946 | |
|  | drt + hwave + veg + bio17 + bio8 + bio9 |  | 0.879 | | 0.792 | | 0.807 | |  | |  | | 0.652 | |  | | 0.782 | |
| Common blossom bat  (*Syconycteris australis*) | drt + hwave + veg + bio14 + bio18 + bio8 | AUC | 0.938 | | 0.822 | | 0.918 | | 0.933 | | 0.880 | | 0.882 | | 0.877 | | 0.893 | |
|  | drt + hwave + veg + bio14 + bio18 + bio9 |  | 0.942 | | 0.820 | | 0.906 | | 0.941 | | 0.888 | | 0.889 | | 0.874 | | 0.894 | |
|  | drt + hwave + veg + bio14 + bio8 + bio9 |  | 0.934 | | 0.732 | | 0.932 | | 0.931 | | 0.880 | | 0.880 | | 0.889 | | 0.883 | |
|  | drt + hwave + veg + bio18 + bio8 + bio9 |  | 0.937 | | 0.821 | | 0.897 | | 0.935 | | 0.858 | | 0.861 | | 0.907 | | 0.888 | |
|  | drt + hwave + veg + bio14 + bio18 + bio8 | TSS | 0.432 | | 0.394 | | 0.396 | | 0.510 | | 0.124 | | 0.118 | | 0.407 | | 0.340 | |
|  | drt + hwave + veg + bio14 + bio18 + bio9 |  | 0.427 | | 0.423 | | 0.296 | | 0.517 | | 0.109 | | 0.102 | | 0.447 | | 0.332 | |
|  | drt + hwave + veg + bio14 + bio8 + bio9 |  | 0.417 | | 0.356 | | 0.486 | | 0.437 | | 0.119 | | 0.100 | | 0.461 | | 0.340 | |
|  | drt + hwave + veg + bio18 + bio8 + bio9 |  | 0.381 | | 0.399 | | 0.349 | | 0.457 | | 0.084 | | 0.086 | | 0.470 | | 0.318 | |
| Northern blossom bat  (*Macroglossus minimus*) | drt + hwave + veg + bio18 + bio5 + bio8 | AUC | 0.976 | | 0.883 | | 0.937 | |  | |  | | 0.945 | |  | | 0.935 | |
|  | **drt + hwave + veg + bio18 + bio5 + bio9** |  | **0.971** | | **0.889** | | **0.958** | |  | |  | | **0.949** | |  | | **0.942** | |
|  | drt + hwave + veg + bio18 + bio8 + bio9 |  | 0.978 | | 0.909 | | 0.944 | |  | |  | | 0.941 | |  | | 0.943 | |
|  | drt + hwave + veg + bio5 + bio8 + bio9 |  | 0.973 | | 0.922 | | 0.968 | |  | |  | | 0.954 | |  | | 0.954 | |
|  | drt + hwave + veg + bio18 + bio5 + bio8 | TSS | 0.621 | | 0.713 | | 0.436 | |  | |  | | 0.599 | |  | | 0.592 | |
|  | **drt + hwave + veg + bio18 + bio5 + bio9** |  | **0.631** | | **0.710** | | **0.648** | |  | |  | | **0.654** | |  | | **0.661** | |
|  | drt + hwave + veg + bio18 + bio8 + bio9 |  | 0.656 | | 0.725 | | 0.596 | |  | |  | | 0.491 | |  | | 0.617 | |
|  | drt + hwave + veg + bio5 + bio8 + bio9 |  | 0.601 | | 0.742 | | 0.601 | |  | |  | | 0.507 | |  | | 0.613 | |
| Eastern tube-nosed bat  (*Nyctimene robinsoni*) | drt + hwave + veg + bio18 + bio19 + bio8 | AUC | 0.888 | | 0.771 | | 0.892 | |  | |  | | 0.896 | |  | | 0.862 | |
|  | drt + hwave + veg + bio18 + bio19 + bio9 |  | 0.895 | | 0.759 | | 0.896 | |  | |  | | 0.903 | |  | | 0.863 | |
|  | drt + hwave + veg + bio18 + bio8 + bio9 |  | 0.900 | | 0.778 | | 0.904 | |  | |  | | 0.900 | |  | | 0.870 | |
|  | drt + hwave + veg + bio19 + bio8 + bio9 |  | 0.852 | | 0.736 | | 0.801 | |  | |  | | 0.866 | |  | | 0.814 | |
|  | drt + hwave + veg + bio18 + bio19 + bio8 | TSS | 0.098 | | 0.485 | | 0.377 | |  | |  | | 0.271 | |  | | 0.308 | |
|  | drt + hwave + veg + bio18 + bio19 + bio9 |  | | 0.082 | | 0.479 | | 0.290 | |  | |  | | 0.299 | |  | | 0.287 |
|  | drt + hwave + veg + bio18 + bio8 + bio9 |  |  | 0.097 | | 0.516 | | 0.380 | |  | |  | | 0.260 | |  | | 0.313 |
|  | drt + hwave + veg + bio19 + bio8 + bio9 |  |  | 0.000 | | 0.161 | | 0.071 | |  | |  | | 0.278 | |  | | 0.128 |

* Predictors: drt = drought, hwave = heatwave, veg= vegetation, bio5 = max temperature of warmest month, bio8 = mean temperature of wettest quarter, bio9 = mean temperature of driest quarter, bio17 = precipitation of driest quarter, bio18 = precipitation of warmest quarter, bio19 = precipitation of coldest quarter

** Accuracy metrics: Area Under the Receiver Operating Characteristic Curve (AUC) and True Skill Statistic (TSS)

*** Machine learning algorithms: random forest (RF), classification and regression trees (CART), neural network (NN), stochastic gradient boosting (GB), penalised generalized linear model (GLM), penalized multinomial regression (MR), flexible discriminant analysis (FDA). Algorithms which failed to fit have blank values.

**Table S1.4.** Cross-validated variable importance of the overall top 10 predictors for the three species for which the models achieved high descriptive accuracy.

| **Species** | **No.** | **Predictor*** | **overall** | **Machine learning algorithm**** | | | | | |
| --- | --- | --- | --- | --- | --- | --- | --- | --- | --- |
|  |  |  |  | **RF** | **CART** | **NN** | **GLM** | **MR** | **FDA** |
| Grey-headed flying fox | 1 | bio8 | 23.937 | 34.336 | 27.682 | 16.503 | 0.390 | 0.400 | 59.688 |
|  | 2 | bio19 | 17.664 | 26.280 | 25.718 | 12.566 | 0.000 | 0.000 | 16.532 |
|  | 3 | bio9 | 15.095 | 19.725 | 21.844 | 17.913 | 0.806 | 0.830 | 11.488 |
|  | 4 | drt | 9.380 | 10.962 | 18.182 | 7.084 | 6.212 | 6.435 | 2.385 |
|  | 5 | hwave | 5.722 | 5.962 | 6.004 | 9.172 | 1.722 | 1.801 | 9.907 |
|  | 6 | veg25 | 3.910 | 1.215 | 0.569 | 8.431 | 7.807 | 8.460 | 0.000 |
|  | 7 | veg18 | 3.902 | 0.007 | 0.000 | 0.998 | 16.809 | 14.447 | 0.000 |
|  | 8 | veg14 | 3.877 | 0.211 | 0.000 | 1.095 | 16.149 | 12.323 | 0.000 |
|  | 9 | veg24 | 3.675 | 0.119 | 0.000 | 2.785 | 14.048 | 14.868 | 0.000 |
|  | 10 | veg2 | 3.389 | 0.744 | 0.000 | 7.518 | 7.436 | 8.082 | 0.000 |
| Spectacled flying fox | 1 | bio18 | 71.632 | 97.300 | 74.127 | 22.917 |  | 1.381 |  |
|  | 2 | bio8 | 12.840 | 0.574 | 12.133 | 37.753 |  | 38.791 |  |
|  | 3 | temp42 | 8.399 | 0.000 | 11.200 | 1.622 |  | 0.000 |  |
|  | 4 | bio17 | 2.246 | 0.943 | 1.331 | 24.439 |  | 1.921 |  |
|  | 5 | drt | 2.094 | 1.062 | 1.208 | 5.828 |  | 14.027 |  |
|  | 6 | veg25 | 1.595 | 0.026 | 0.000 | 3.725 |  | 25.605 |  |
|  | 7 | veg2 | 1.064 | 0.095 | 0.000 | 0.530 |  | 18.273 |  |
|  | 8 | veg24 | 0.042 | 0.000 | 0.000 | 1.032 |  | 0.000 |  |
|  | 9 | veg14 | 0.034 | 0.000 | 0.000 | 0.843 |  | 0.000 |  |
|  | 10 | veg19 | 0.017 | 0.000 | 0.000 | 0.407 |  | 0.000 |  |
| Northern blossom bat | 1 | bio9 | 24.465 | 31.238 | 33.053 | 8.893 |  | 0.516 |  |
|  | 2 | bio18 | 17.685 | 14.003 | 4.700 | 46.491 |  | 0.000 |  |
|  | 3 | bio5 | 17.673 | 20.253 | 19.026 | 16.544 |  | 1.174 |  |
|  | 4 | drt | 16.262 | 19.242 | 23.323 | 6.216 |  | 4.636 |  |
|  | 5 | hwave | 8.807 | 6.320 | 7.883 | 3.836 |  | 40.145 |  |
|  | 6 | veg2 | 5.313 | 4.670 | 10.480 | 1.439 |  | 4.305 |  |
|  | 7 | veg19 | 2.199 | 1.408 | 1.534 | 2.378 |  | 8.471 |  |
|  | 8 | veg6 | 1.959 | 0.786 | 0.000 | 3.386 |  | 10.981 |  |
|  | 9 | veg25 | 1.926 | 1.265 | 0.000 | 4.739 |  | 3.980 |  |
|  | 10 | veg7 | 1.329 | 0.591 | 0.000 | 2.103 |  | 7.633 |  |

* Predictors: drt = drought, hwave = heatwave, bio5 = max temperature of warmest month, bio8 = mean temperature of wettest quarter, bio9 = mean temperature of driest quarter, bio17 = precipitation of driest quarter, bio18 = precipitation of warmest quarter, bio19 = precipitation of coldest quarter, veg25= cleared, non-native vegetation, buildings, veg18= heathlands, veg14= mallee, veg24= inland Aquatic and mangroves, veg2= Eucalypt, veg19= grasslands, veg6= Acacia, veg7= other forests and woodlands.

** Machine learning algorithms: random forest (RF), classification and regression trees (CART), neural network (NN), generalized linear model (GLM), penalized multinomial regression (MR), flexible discriminant analysis (FDA). Algorithms which were not used to fit have blank values.

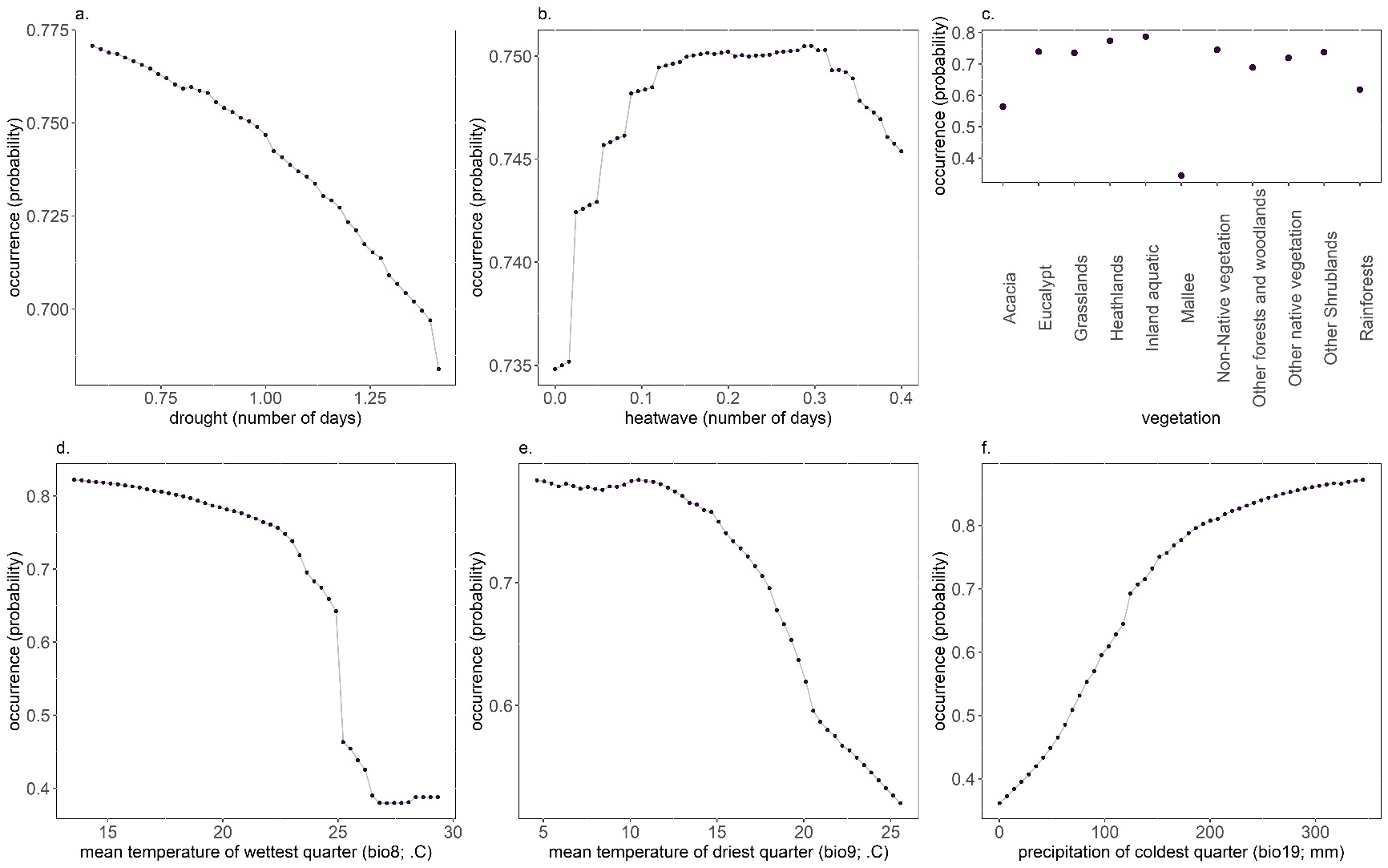

**Figure S1.1**. Partial dependence plot of the drought (a), heatwave (b), vegetation (c), mean temperature of wettest quarter (d), mean temperature of driest quarter (e) and precipitation of coldest quarter (f) on the probability of occurrence of the Grey-headed flying fox.

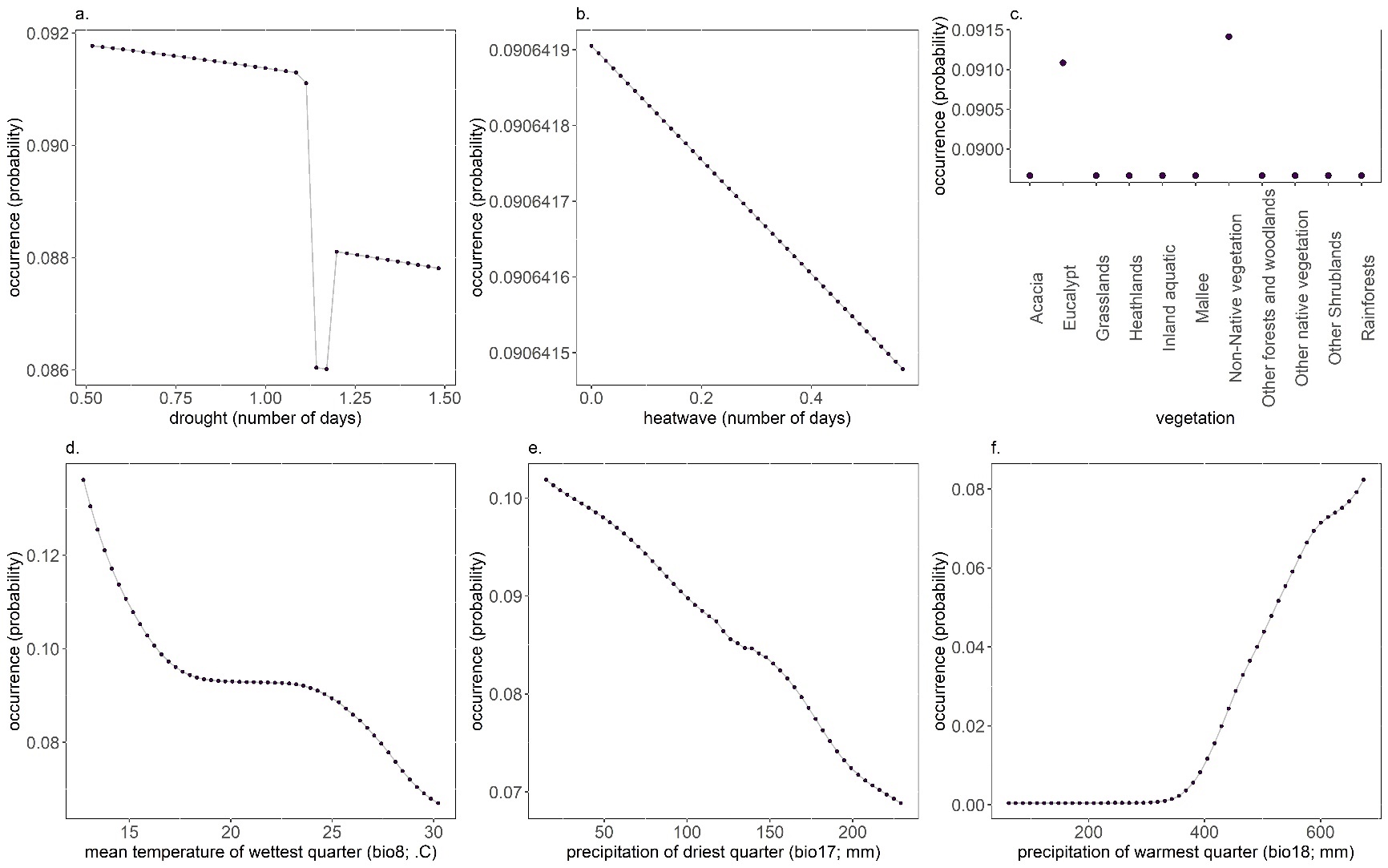

**Figure S1.2** Partial dependence plot of the drought (a), heatwave (b), vegetation (c), mean temperature of wettest quarter (d), precipitation of driest quarter (e) and precipitation of warmest quarter (f) on the probability of occurrence of the Spectacled flying fox.

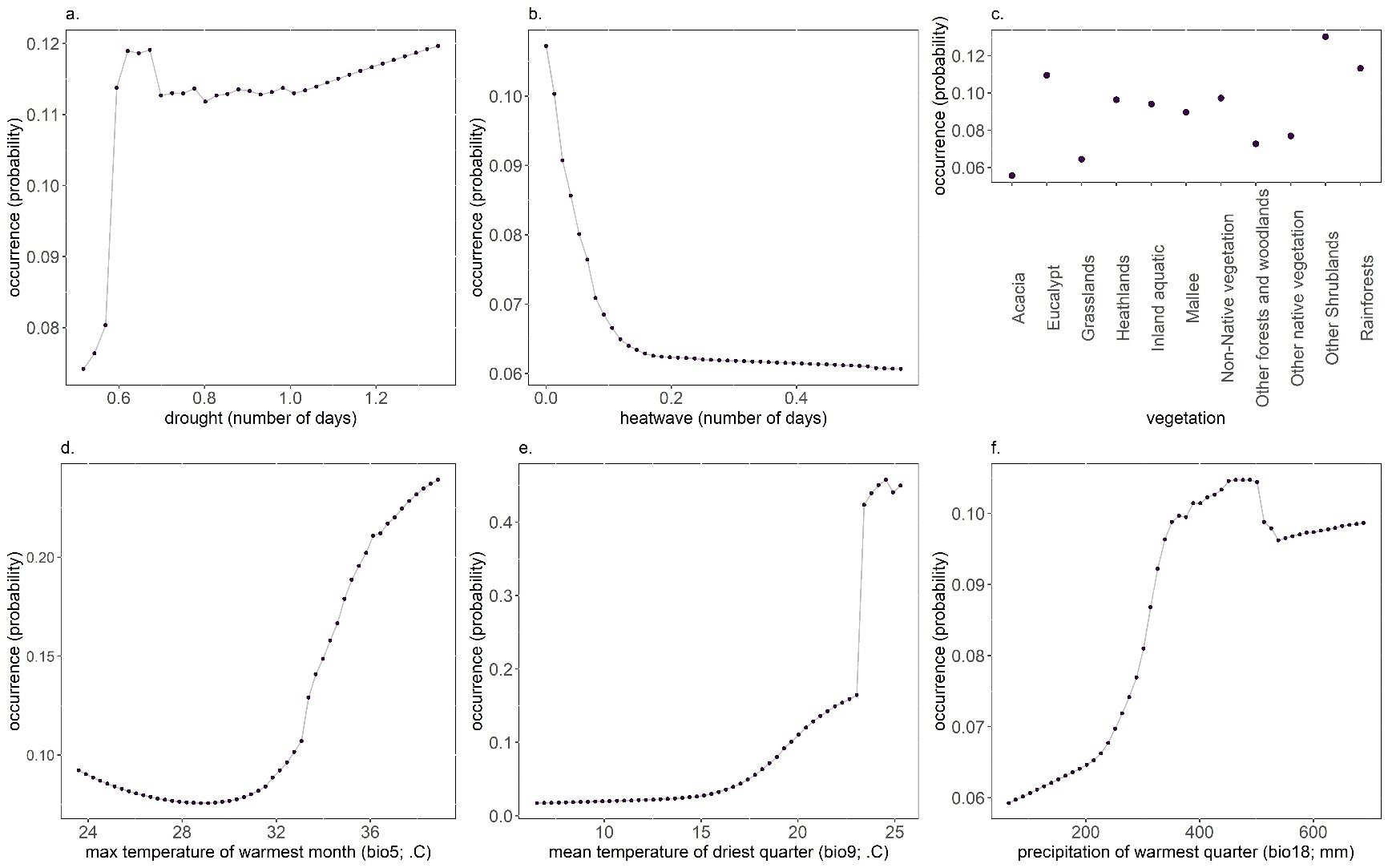

**Figure S1.3** Partial dependence plot of the drought (a), heatwave (b), vegetation (c), max temperature of wettest month (d), mean temperature of driest quarter (e) and precipitation of warmest quarter (f) on the probability of occurrence of the Northern blossom bat.

**Table S1.5** Percentage of areas suitable, unsuitable, lost and gain under different future scenarios for each species.

| **Species** | **Areas** | **2050** | |  | **2070** | |
| --- | --- | --- | --- | --- | --- | --- |
|  |  | **rcp 4.5** | **rcp 8.5** |  | **rcp 4.5** | **rcp 8.5** |
| Grey-headed flying fox | Suitable | 10.049 | 7.906 |  | 9.352 | 6.709 |
|  | Unsuitable | 83.624 | 83.624 |  | 83.540 | 83.418 |
|  | Lost | 5.662 | 7.805 |  | 6.358 | 9.001 |
|  | Gained | 0.663 | 0.663 |  | 0.748 | 0.870 |
| Spectacled flying fox | Suitable | 0.270 | 0.196 |  | 0.221 | 0.291 |
|  | Unsuitable | 99.293 | 99.408 |  | 99.374 | 99.380 |
|  | Lost | 0.185 | 0.259 |  | 0.235 | 0.164 |
|  | Gained | 0.250 | 0.135 |  | 0.168 | 0.163 |
| Northern blossom bat | Suitable | 3.567 | 3.497 |  | 3.941 | 3.559 |
|  | Unsuitable | 92.312 | 92.214 |  | 92.043 | 91.763 |
|  | Lost | 2.112 | 2.181 |  | 1.738 | 2.120 |
|  | Gained | 2.007 | 2.106 |  | 2.277 | 2.557 |

**Table S1.6** Percentage of presence of Grey-headed flying fox, Spectacled flying fox, and Northern blossom bat across different climate types under current conditions (when compared to present climate type based on Beck et al. 2018) and future scenarios (when compared to future climate type based on Beck et al. 2018). Climate description area based on Köppen-Geiger climate classification see Beck et al. 2018.

|  |  |  |  | **2050** | |  | **2070** | |
| --- | --- | --- | --- | --- | --- | --- | --- | --- |
| **Species** | **Climate code** | **Climate type** | **current** | **rcp 4.5** | **rcp 8.5** |  | **rcp 4.5** | **rcp 8.5** |
| Grey-headed flying fox | 1 | Tropical, rainforest |  |  |  |  |  |  |
|  | 2 | Tropical, monsoon | 0.021 |  |  |  |  |  |
|  | 3 | Tropical, savannah |  | 0.013 |  |  | 0.007 |  |
|  | 4 | Arid, desert, hot |  |  |  |  |  |  |
|  | 5 | Arid, desert, cold |  |  |  |  |  |  |
|  | 6 | Arid, steppe, hot |  | 0.371 | 0.063 |  | 0.586 | 0.027 |
|  | 7 | Arid, steppe, cold | 10.821 | 7.857 | 5.519 |  | 8.229 | 5.040 |
|  | 8 | Temperate, dry summer, hot summer | 7.833 | 8.265 | 2.771 |  | 6.030 | 1.100 |
|  | 9 | Temperate, dry summer, warm summer | 17.042 | 9.559 | 9.444 |  | 9.575 | 10.453 |
|  | 11 | Temperate, dry winter, hot summer | 0.201 |  |  |  |  |  |
|  | 14 | Temperate, no dry season, hot summer | 21.446 | 45.111 | 46.075 |  | 44.276 | 39.423 |
|  | 15 | Temperate, no dry season, warm summer | 42.623 | 28.826 | 36.128 |  | 31.297 | 43.957 |
|  | 16 | Temperate, no dry season, cold summer |  |  |  |  |  |  |
|  | 26 | Cold, no dry season, warm summer | 0.013 |  |  |  |  |  |
|  | 27 | Cold, no dry season, cold summer |  |  |  |  |  |  |
|  | 29 | Polar, tundra |  |  |  |  |  |  |
|  |  |  |  | **2050** | |  | **2070** | |
| **Species** | **Climate code** | **Climate type** | **current** | **rcp 4.5** | **rcp 8.5** |  | **rcp 4.5** | **rcp 8.5** |
| Spectacled flying fox | 1 | Tropical, rainforest | 10.029 | 6.839 | 10.751 |  | 9.138 | 7.829 |
|  | 2 | Tropical, monsoon | 25.811 | 29.032 | 40.568 |  | 33.621 | 33.235 |
|  | 3 | Tropical, savannah | 34.956 | 62.839 | 48.276 |  | 56.379 | 58.936 |
|  | 4 | Arid, desert, hot |  |  |  |  |  |  |
|  | 5 | Arid, desert, cold |  |  |  |  |  |  |
|  | 6 | Arid, steppe, hot |  |  |  |  |  |  |
|  | 7 | Arid, steppe, cold |  |  |  |  |  |  |
|  | 8 | Temperate, dry summer, hot summer |  |  |  |  |  |  |
|  | 9 | Temperate, dry summer, warm summer |  |  |  |  |  |  |
|  | 11 | Temperate, dry winter, hot summer | 27.434 |  |  |  |  |  |
|  | 14 | Temperate, no dry season, hot summer | 1.770 | 1.290 | 0.406 |  | 0.862 |  |
|  | 15 | Temperate, no dry season, warm summer |  |  |  |  |  |  |
|  | 16 | Temperate, no dry season, cold summer |  |  |  |  |  |  |
|  | 26 | Cold, no dry season, warm summer |  |  |  |  |  |  |
|  | 27 | Cold, no dry season, cold summer |  |  |  |  |  |  |
|  | 29 | Polar, tundra |  |  |  |  |  |  |
|  |  |  |  | **2050** | |  | **2070** | |
| **Species** | **Climate code** | **Climate type** | **current** | **rcp 4.5** | **rcp 8.5** |  | **rcp 4.5** | **rcp 8.5** |
| Norther blossom bat | 1 | Tropical, rainforest | 0.024 | 0.339 | 0.637 |  | 0.574 | 0.583 |
|  | 2 | Tropical, monsoon | 0.059 | 0.980 | 2.694 |  | 2.059 | 2.576 |
|  | 3 | Tropical, savannah | 99.560 | 98.620 | 96.537 |  | 97.313 | 96.654 |
|  | 4 | Arid, desert, hot |  |  |  |  |  |  |
|  | 5 | Arid, desert, cold |  |  |  |  |  |  |
|  | 6 | Arid, steppe, hot | 0.356 | 0.048 | 0.012 |  | 0.033 | 0.066 |
|  | 7 | Arid, steppe, cold |  |  |  |  |  |  |
|  | 8 | Temperate, dry summer, hot summer |  | 0.012 | 0.120 |  | 0.022 | 0.121 |
|  | 9 | Temperate, dry summer, warm summer |  |  |  |  |  |  |
|  | 11 | Temperate, dry winter, hot summer |  |  |  |  |  |  |
|  | 14 | Temperate, no dry season, hot summer |  |  |  |  |  |  |
|  | 15 | Temperate, no dry season, warm summer |  |  |  |  |  |  |
|  | 16 | Temperate, no dry season, cold summer |  |  |  |  |  |  |
|  | 26 | Cold, no dry season, warm summer |  |  |  |  |  |  |
|  | 27 | Cold, no dry season, cold summer |  |  |  |  |  |  |
|  | 29 | Polar, tundra |  |  |  |  |  |  |
