## Appendix S2 for "Predicted the impacts of climate change and extreme-weather events on the future distribution of fruit bats in Australia"

**Table S2.1** Overview, Data, Model, Assessment and Prediction (ODMAP) protocol

| **OVERVIEW** | |
| --- | --- |
| Authorship | - Title: Predicted the impacts of climate change and extreme-weather events on the future distribution of fruit bats in Australia - Authors: Vishesh L. Diengdoh*, Stefania Ondei, Mark Hunt and Barry W. Brook - Contact email: * |
| Model objective | - SDM objectives: predict occurrence under current conditions and future scenarios. - Target outputs: probability of occurrence maps and binary maps representing presence and absence. |
| Taxon | Fruit bats (Megachiroptera) |
| Location | Australia |
| Scale of analysis | - Spatial extent: 112.925, 155.525, -43.875, -9.975 (xmin, xmax, ymin, ymax) - Spatial resolution: 0.05° (∼ 5 km) - Temporal extent: Current (2020), Future (2050 and 2070) - Type of extent boundary: Political (Australia) |
| Biodiversity data | - Observation type: field survey (human observation) - Response/data type: presence (field data) and pseudo absence (generated) |
| Type of predictors | drought, heatwaves, vegetation, bioclimatic |
| Conceptual model | Fruit bats (Megachiroptera) are threatened by changes in climate and sensitive to extreme-weather events. We assessed the potential impacts of those changes, particularly more frequent and intense heatwaves and drought, on the future distribution of fruit bats in Australia. Correlative species distribution models (SDM) have been used, to assess the impacts of climate change on fruit bats, but either failed to account for changes in extreme weather or did not use species-specific temperature thresholds. |
| Assumptions | - Heatwaves (>=42 °C) is a temperature threshold for all study species. - Vegetation is a constant predictor under current and future scenarios. - Relevant predictors for fruit bat distributions are included. - Study area excluded desert regions because it is unlikely that these regions will be suitable for fruit bats due to their high temperatures and extreme range. - Bias in presence and pseudo-absence have been accounted. |
| SDM algorithms | - Algorithms: Random forest (RF), classification and regression trees (CART), neural network (NN), stochastic gradient boosting (GB), penalised generalized linear model (GLM), penalised multinomial regression (MR) and flexible discriminant analysis (FDA). - Ensemble: unweighted averaging of the algorithms. |
| Model workflow | The training data was biased to pseudo-absences (1 presence to 10 pseudo-absence). Pseudo-absences were selected using a stratified random method to ensure adequate number vegetation class (the only factorial predictor) were represented.  Correlated predictors were removed using a threshold of 0.7. If two variables have a high correlation, the variable with the largest mean absolute correlation is removed. Instead of using all the non-correlated predictors to fit the models, we created a list of all possible combinations of three non-correlated bioclimatic predictors and included heatwave, drought, and vegetation predictors since they were not strongly correlated to each other or any of the climate variables. This resulted in a set of four different group of predictors/candidate models for each species.  We used repeated 70/30% training/test cross-validation splits with 50 repeats for algorithm tuning/optimisation, selecting of predictor/candidate models for each species, and calculating variable importance.  Occurrence models were then re-fitted to the full dataset using the best predictor/candidate models derived from the cross-validation step. We also assessed the partial dependence plots of each algorithm and then averaged them (unweighted) per best predictor/candidate model. We then averaged the algorithms using an unweighted method. Future-occurrence predictions were made for each species and then ensembled (unweighted) per year per emission scenario for each species.  We transformed the probabilistic ensemble occurrence models to binary (presence and absence representations) using a sensitivity‐specificity sum maximisation approach to select the optimal suitability threshold. |
| Software | - Modelling platform: R software (R Core Team 2020) - Data availability: training data for each study species, and occurrence and binary (presence/absence) maps (under current conditions and future scenarios as GeoTiff) for only the Grey-headed flying fox, Spectacled flying fox, and Northern blossom bat have been made publicly available. - Code availability: code not included because we used generic functions provided in the different r packages. |
| **DATA** | |
| Biodiversity data | - Taxon names: Grey-headed flying-fox (*Pteropus poliocephalus*), Little red flying-fox (*P. scapulatus*), Black flying-fox (*P. alecto*), Spectacled flying-fox (*P. conspicillatus*), Common blossom bat (*Syconycteris australis*), Northern blossom bat (*Macroglossus minimus*), and Eastern tube-nosed bat (*Nyctimene robinsoni*) - Ecological level: species level - Data source: Atlas of living Australia (ALA) - Sampling design: unknown - Sample size (for training data): Grey-headed flying fox (0= 713, 1=1940), Little red flying fox (0=1911, 1=742), Black flying fox (0=1943, 1=710), Spectacled flying fox (0=240, 1=24), Common blossom bat (0=440, 1=44), Northern blossom bat (0=240, 1=24), Eastern tube-nosed bat (0=280, 1=28). Here, 0= pseudo-absence and 1= presence. - Regional mask: Australian continent, with the exclusion of desert regions, as identified using a Köppen-Geiger climatic layer including the land-cover categories: extraction sites, salt lakes and alpine grassland. - Scaling: Presence data was reduced to a single observation per grid cell of 0.05° (∼ 5 km). - Data cleaning: Occurrence records were limited to those from 1960 and onwards and classified as ‘human observation’. Duplicate records based on latitude and longitude and dubious records (e.g., outliers well outside the known distribution ranges) were removed. - Background data/pseudo-absences: pseudo-absences were generated using the target‐group method. The method consists in generating pseudo-absences for each species by using the presence points of other fruit bats. - Errors and biases: Pseudo-absences were generated using target‐group method as it can handle sampling bias. If the presence and pseudo-absence data have the same bias, the model will ignore the sample selection bias and focus on other differentiation between the distribution of the presences and pseudo-absence data. Apart from this we did not account/correct for any other biases. |
| Data partitioning | No data partitioning was applied. Model performance was assessed using repeated 70/30% training/test cross-validation splits with 50 repeats. Ensemble model performance was assessed using the entire training data i.e., goodness of fit. |
| Predictor variables | - Predictor variables: heatwave (the number of days where temperature is equal to or greater than 42 °C), drought (the number of months falling below the historic 10^th^ percentile rainfall), vegetation (Appendix S1 Table S1.1), bioclimatic (Appendix S1 Table S1.2). - Data sources: heatwave and drought (Climate Change in Australia, <https://www.climatechangeinaustralia.gov.au>), vegetation (v 5.1, Australian Government Department of Agriculture Water and the Environment, <https://www.environment.gov.au/land/native-vegetation/national-vegetation-information-system/data-products#mvg51>), bioclimatic (current, v 2.1, <https://www.worldclim.org> and future, v 1.4, <http://worldclim.com/> ) - Spatial resolution: 0.05° (∼ 5 km) - Spatial extent: 112.925, 155.525, -43.875, -9.975 (xmin, xmax, ymin, ymax) - Coordinate Reference Systems: +proj=longlat +datum=WGS84 +no_defs - Temporal resolution: Current (2020), Future (2050 and 2070) - Data processing: bioclimatic variables and vegetation were resampled using bilinear interpolation and nearest neighbour method, respectively, to match the spatial resolution of the heatwave and drought predictors. |
| **MODEL** | |
| Variable pre-selection | Drought, heatwaves, vegetation and 12 bioclimatic variables (Appendix S1 Table S1.2) were considered important for fruit bats. |
| Multicollinearity | Correlated predictors were removed using a threshold of 0.7. If two variables were highly correlated, the variable with the largest mean absolute correlation was removed. Instead of using all the non-correlated predictors to fit the models, we created a list of all possible combinations of three non-correlated bioclimatic predictors and included heatwave, drought, and vegetation predictors since they were not strongly correlated to each other or any of the climate variables. This resulted in a set of four different group of predictors/candidate models for each species (see Appendix S1 Table S1.3). |
| Model settings | - Grey-headed flying fox: RF mtry = 5; CART cp = 0.002244039; NN size = 9 and decay = 0.1; GLM alpha = 0.775 and lambda = 0.0002563009; MR decay = 0.01; FDA degree = 1 and nprune = 16. - Spectacled flying fox: RF mtry = 15; CART cp = 0.71875; NN size = 9 and decay = 0.01; MR decay = 0.1. - Northern blossom bat: RF mtry = 5; CART cp = 0.15625; NN size = 9 and decay = 0.01; MR decay = 0.001.   Here: RF= Random forest, CART= classification and regression trees (CART), NN= neural network, GLM= penalised generalized linear model, MR= penalized multinomial regression and FDA= flexible discriminant analysis.  We used the *caret* and *caretEnsemble* R packages to perform repeated 70/30% training/test cross-validation splits with 50 repeats for algorithm tuning. The largest value of AUC was used to select the optimal hyperparameter.  Model settings are only provided for the Grey-headed flying fox (based on the best predictors/candidate models i.e., drt + hwave + veg + bio19 + bio8 + bio9), Spectacled flying fox (drt + hwave + veg + bio18 + bio17 + bio8) and Northern blossom bat (drt + hwave + veg + bio18 + bio5 + bio9).  Model setting are not provided for the Little red flying fox, Black flying fox, Eastern tube-nosed, and Common blossom bat as we did not predict the occurrence for these species due to low cross-validation accuracy (see Appendix S1 Table S1.3).  Here: drt = drought, hwave = heatwave, veg= vegetation, bio5 = max temperature of warmest month, bio8 = mean temperature of wettest quarter, bio9 = mean temperature of driest quarter, bio18 = precipitation of warmest quarter, bio17 = precipitation of driest quarter, bio19 = precipitation of coldest quarter. |
| Model estimates | Assessment of variable importance: variable importance for each algorithm and overall importance were assessed during cross-validation for each species based on the best predictors/candidate models (Appendix S1 Table S1.4) using the caret r package. |
| Model selection/ Ensembles | - Predictors/candidate model selection: the best predictor/candidate models were derived from the cross-validation step. We calculated the AUC and TSS scores of each algorithm and then averaged (unweighted) them (Appendix S1 Table S1.3). The model with the higher TSS and AUC score was considered as the best model. - Ensemble: unweighted averaging of the algorithms. |
| Non-independence correction | None |
| Threshold selection | We used the sensitivity‐specificity sum maximisation approach to select the optimal suitability threshold for transforming the probabilistic ensemble occurrence models to binary (presence and absence representations). |
| **ASSESSMENT** | |
| Performance statistics | - Cross-validation: AUC and TSS (repeated 70/30% training/test cross-validation splits with 50 repeats) - Ensemble model validation: AUS and TSS (goodness of fit) |
| Plausibility checks | None |
| **PREDICTION** | |
| Prediction output | Probabilistic occurrence models and binary (presence and absence) models. We transformed the probabilistic ensemble occurrence models to binary (presence and absence) using a sensitivity‐specificity sum maximisation approach to select the optimal suitability threshold. |
| Uncertainty quantification | None |
